# Selection favors context-dependent bias in altruism

**DOI:** 10.64898/2026.04.26.720906

**Authors:** Linnéa M Båvik, Rohan S Mehta, Daniel B Weissman

**Affiliations:** Department of Physics, Emory University; School of Biological Sciences, Georgia Institute of Technology; Department of Biology, Elmhurst University

**Keywords:** altruism, cooperation, evolutionary dynamics, evolutionary game theory, mathematical biology

## Abstract

Altruism, in which individuals sacrifice some of their own reproduction to help others, can evolve if it is preferentially directed toward relatives. Organisms may recognize relatives through phenotypic similarity. Under the models originally studied by Hamilton, the threshold relatedness at which altruism becomes beneficial depends on the overall relatedness of the population. Implicit in this result is that natural selection may favor context-dependent strategies, in which donors judge their similarity to potential recipients relative to their similarity to the overall population when deciding whether to help. In this manuscript, we use a combination of simulations and theory to determine the circumstances under which context dependence is favored over simple strategies that do not depend on the observation of other interacting individuals. We find that a “plastic” strategy that uses the population context in its rule for donating consistently beats strategies that use only information about the potential recipient.

## Introduction

One lesser-discussed aspect of Hamilton’s rule for the evolution of social traits (*r > c/b*) is that relatedness is not only a function of the genetic distance between an actor and recipient, but a function of their population context as well. As Hamilton himself wrote, the “omniscient animal” would be at best “indifferent to any individuals having the average relationship” [Hamilton, 1970]. In other words, the crucial question is how closely the recipient is related to the actor *compared to others individuals in the population*, not on an absolute scale. This point is often overlooked in popular descriptions of altruism, such as Haldane’s supposed quip that he was “prepared to lay down his life for eight cousins or two brothers”, which implicitly assumes that the average individual in the population is much less related to him than his cousins are [Maynard Smith, 1975, O’Toole, 2017].

Many theoretical papers have emphasized that relatedness depends on population composition [Hamilton, 1975, Grafen, 1985, Queller, 1994, Taylor, 1992, Gardner and West, 2004, Krupp and Taylor, 2013, 2015, Madgwick, 2020], and thus population composition can drive or reverse the inclusive fitness incentives for altruism. For example, spatial structure can induce competition among relatives that mitigates kin selection for altruism. But given that social behavior often involves sensing the environment before taking action [Allen et al., 2016], less attention has been paid to how population composition observed in the sensing stage could then be utilized in the individual’s decision making. Particularly when the genetic environment of a parent and offspring may vary, the ability to modulate behavior according to true relatedness of one’s partners allows individuals to tailor their helping behaviors to be more generous in environments when the altruism gene is scarce and stingier when the gene is close to fixed, as incurring costs to help the “average” individual in this population has no effect on increasing gene frequency and thus by definition removes the selective advantage of carrying the gene.

In order to understand optimal discrimination among kin, it is necessary first to understand how individuals infer kinship. One well-supported mechanism for kin recognition is called phenotype matching: individuals observe the phenotype of a partner in order to gain information about the underlying genotype [Waldman, 1987, Krupp et al., 2011]. This mechanism underlies a host of theories for the evolution of social traits including the “greenbeard effect” [Dawkins, 1976], the “armpit effect” [Dawkins, 1982], “tag-based cooperation” [Riolo et al., 2001, Axelrod et al., 2004], and models of “conspecific acceptance thresholds” [Reeve, 1989]. Phenotype matching has been widely observed in nature and experiments in taxa ranging from insects, birds, and mammals to plants [Scharf et al., 2020]. However, there has been little study of if or how individuals sense the composition of the background population.

Recent work has highlighted the potential relevance of sensing population composition to social behavior involving kin recognition. On the theory side, Krupp and Taylor [2013] pointed out that phenotype similarity can only be used to infer genotypic relatedness when compared to the population phenotype distribution: in their model, the potential recipient only has high genetic relatedness when their phenotype is closer to that of the altruist than the population average and the altruist-recipient phenotypic distance is small relative to the underlying standard deviation of the phenotype distribution of the population as a whole. In addition, in species where dispersal is common, individuals can frequently find themselves in environments of asymmetrical relatedness as the quantity of migrant individuals can be far outnumbered by the population of native individuals. In this case, learning, rather than inheriting, information about the population’s phenotypic distribution allows individual behavior to plastically adhere to the underlying inclusive fitness incentives which have been shown to span the range between altruism and spite as a function of population composition alone [Krupp and Taylor, 2015]. The Trinidadian guppy *Poecilia reticulata* is such a species, as these fish reside in a pool-and-riffle ecology where individuals disperse between semi-isolated pools, and population demography fluctuates with location and season. Strikingly, Daniel [2020] conducted controlled experiments and found that these fish did indeed plastically alter social mating behaviors according to population phenotypic context in agreement with the theory.

Empirical work has found behavior that is dependent on population context in taxa ranging from microbes to insects to vertebrates, like birds and mammals. The altruistic spore-forming amoeba *Dictyostelium discoidium* exhibits population context-dependent behavior in the presence of strains of varying relatedness [Noh et al., 2020]. The Argentine ant *Linepithema humile*, when living in colonies of greater phenotypic variability, exhibits decreased aggression toward conspecifics introduced from foreign lesser related colonies [Tsutsui et al., 2003]. Similarly, the Columbian ground squirrel *Spermophilus columbianus*, when exposed to olfactory cues from neighboring colonies, showed more cohesive behavior toward non-colony members than squirrels reared without these cues [Hare, 1994]. Other organisms that have been noted to use population estimation in social decision making include great reed warblers, weaver ants, and humans ([Moskát et al., 2008, Newey et al., 2010, Krupp et al., 2012], as reviewed by Krupp and Taylor [2013]).

In wild populations, Hamilton’s hypothetical “omniscient animal” that knows its exact relatedness to all potential partners does not exist. Animals have at best limited, noisy data about the composition of their population. This raises the question: when will natural selection favor trying to use this information to decide who to help? To address this question, we develop an evolutionary simulation model in which different genotypes have different strategies for kin recognition and altruistic behavior. One set of “innate” strategies, whose evolution we studied in [Båvik et al., 2023], have a genetically encoded threshold of phenotypic similarity that they use in determining when to behave altruistically. The other set of strategies are “plastic”. They too use a threshold of phenotypic similarity in determining when to behave altruistically, but what is inherited is not the threshold itself but rather a simple rule that converts limited observations of the population diversity into a threshold. In other words, the individuals following an innate strategy are born with an internal scale for measuring phenotypic similarity, while those following a plastic strategy have to learn the scale from their interactions. We find that the strategy most often favored by natural selection is a plastic one that judges potential partners based on population context. However, we also find population configurations where plastic individuals cannot effectively use the limited population information available, and natural selection favors innate strategies.

### Model

Our model focuses on the contrast between innate and plastic strategies for kin recognition (Figure 1). Individuals who have an innate strategy evaluate a potential partner based solely on the partner’s phenotype, irrespective of population context. An innate strategist thus regards a partner as an equally viable recipient of altruism whether the background population is highly homogeneous or extremely heterogeneous. In contrast, individuals who have a plastic strategy evaluate a potential recipient of altruism differently depending on the population context. A plastic strategist will choose whether or not to donate to a recipient based on that recipient’s *relative* similarity compared to the rest of the individuals with which they are interacting.

**Figure 1:**
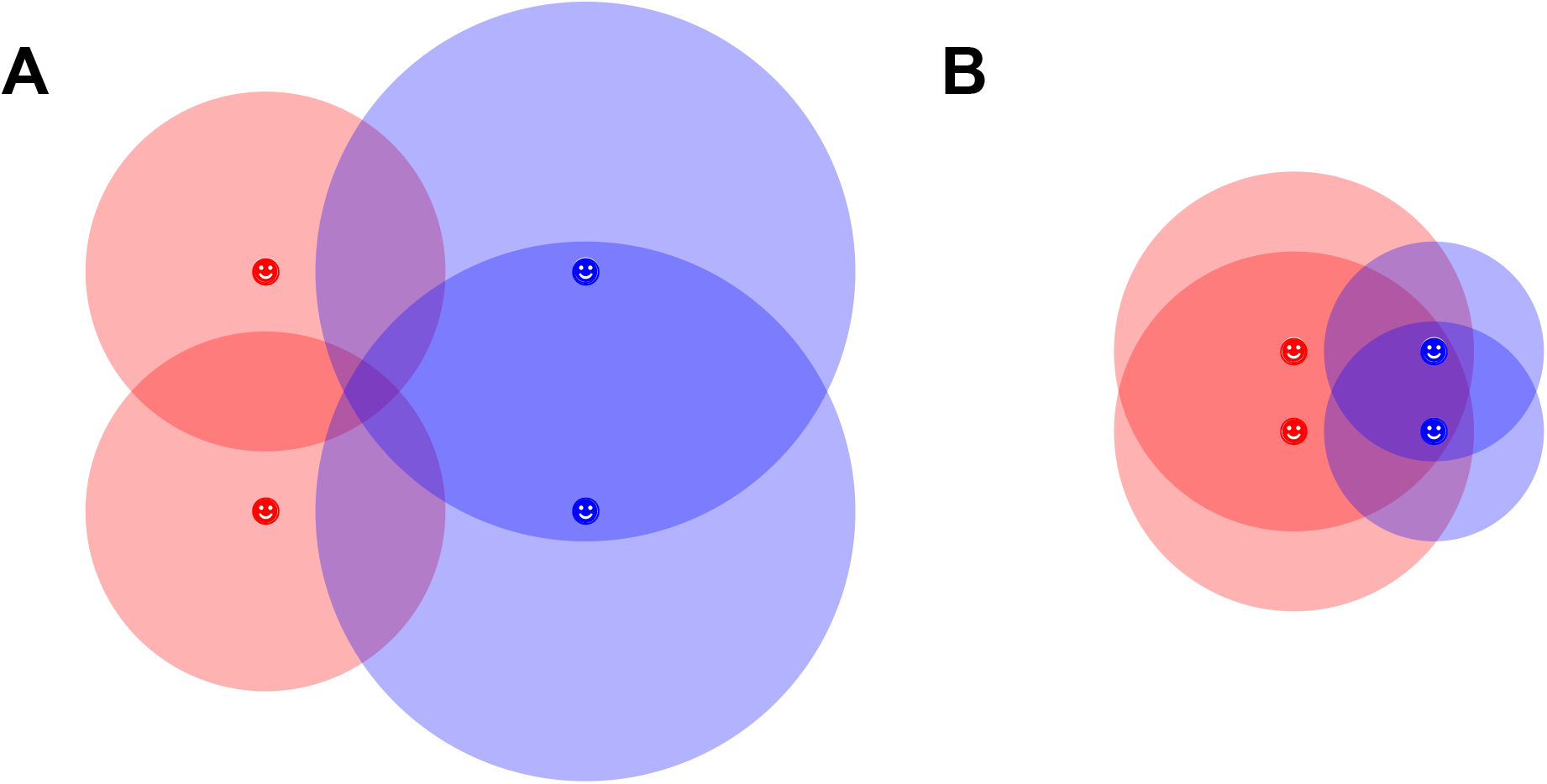
Schematic of altruistic behavior of plastic strategists (blue) and innate strategists (red). Smiley faces indicate the locations of individuals in a two-dimensional phenotype space that represents the components of their phenotype visible to other members of the population. The radius extending from each individual indicates their altruism threshold: partners within the altruism threshold will be the recipients of a donation, those outside will not. This schematic illustrates the advantage of plasticity: whether the population is diffuse (Panel **A**) or compact (Panel **B**) in phenotype space, plastic altruists can flexibly funnel helping towards kin, while innate altruists may fail to help kin in some situations and be easily cheated by nonkin in others.

Our specific model is essentially identical to that of Båvik et al. [2023], with the addition of the plastic strategies; for completeness we will describe it here. All parameter values can be found in Table 1. We model the population as well-mixed and consisting of *N* asexual individuals, where each individual *i* has a phenotype Φ_*i*_ ∈ ℝ ^*D*^—a point in a *D*-dimensional continuous phenotype space that can be observed by other individuals in the population—and an unobservable strategy, which consists of a calculated threshold 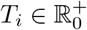 that determines whether or not it will donate to another individual. Each generation consists of two steps: interaction and reproduction. In the interaction step, a focal individual *i* observes a random sample of *K* potential recipients. For each partner *k* in that sample, the actor *i* calculates the Euclidean distance between its own phenotype and the partner’s (*d*_*ij*_ = ||Φ_*i*_ −Φ_*j*_||). Individual *i* compares this distance to their threshold *T*_*i*_, and bestows an altruistic donation on *k* if *d*_*ik*_ *< T*_*i*_. This process corresponds to an organism comparing a conspecific to its kin template during an interaction. If *i* does not donate to *k, i*’s payoff is increased by *c* and *k*’s is unchanged, while if *i* does donate, *i*’s payoff is unchanged and *k*’s is increased by *b*, with *b > c*.

**Table 1:** Model parameters.

| Symbol | Description | Range | Figure 2 | Figure 3 | Figure 4 | Figure 5 |
| --- | --- | --- | --- | --- | --- | --- |
| $N$ | Number of individuals in population | $[1, \infty)$ | 1000 | 1000 | 1000 | 1000 |
| | Number of generations for simulation | $[1, \infty)$ | $1 \times 10^4$ | $1 \times 10^4$ | $5 \times 10^3, 5 \times 10^4$ | $2 \times 10^7, 1 \times 10^5$ |
| $b$ | Benefit of altruism | $[0, \infty)$ | 10 | 10 | 10 | 10 |
| $c$ | Cost of altruism | $[0, \infty)$ | 1 | 1 | 1 | 1 |
| $T$ | Threshold for altruism | $[0, 28.446]$ | N/A | 14.2 | Fixed or N/A | N/A |
| $a$ | Plastic threshold coefficient | $[0, 2]$ | N/A | 0.8 | Fixed or N/A | N/A |
| $K$ | Sample size | $[0, \infty)$ | 10 | 10 | 10 | 10 |
| $s$ | Strength of selection | $[0, \infty)$ | 1 | 1 | 1 | 1 |
| | Standard deviation of phenotype mutation distribution | $[0, \infty)$ | 1 | 1 | 1 | 1 |
| $u_s$ | Probability of intra-strategy mutation | $[0, 1]$ | 0.001 | 0 | 0 or 0.001 | 0.001 |
| $u_S$ | Probability of inter-strategy mutation | $[0, 1]$ | 0.001 | 0.001 | 0 or 0.001 | 0.001 |
| $D$ | Phenotype dimension | $\mathbb{Z}^+$ | 20 | 20 | 20 | 20 |

The only difference between the innate and plastic strategies is in how the threshold *T*_*i*_ is determined. If individual *i* is an innate strategist, *T*_*i*_ is simply inherited from its parent (with possible mutation). If individual *i* is a plastic strategist, rather than inheriting a threshold directly, they inherit a scaling factor *a*_*i*_ from their parent. To obtain their threshold *T*_*i*_, they calculate the average dissimilarity of their sample of *K* potential partners, 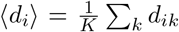, and scale this by *a*_*i*_: *T*_*i*_ = *a*_*i*_ ⟨*d*_*i*_⟩. Larger values of *a*_*i*_ therefore correspond to more altruistic strategies.

The sum of each individual’s interactions of altruistic donations and self-payoffs are accumulated into total payoffs *W* for each individual, where each individual begins with a payoff of 1, such that even the most altruistic possible individual *i* who foregoes all self-payoffs, has at a minimum *W*_*i*_ ≥ 1. An individual’s fitness (expected number of offspring) *w* is proportional to its payoff, normalized by the average payoff 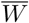 of the population:

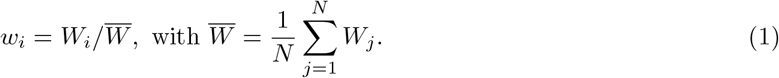

Uniparental reproduction is assumed and offspring are drawn from a multinomial distribution in which the probability of drawing parent *i* is *w*_*i*_*/N*.

Offspring inherit their parent’s phenotype and strategy, plus deviations from mutation. Phenotypes are considered quantitative traits emerging from interactions among many loci and are modeled via the addition of normal noise in every generation. For each dimension of the phenotype, mutations are independently drawn from a standard normal distribution 𝒩 (0, 1) so that the phenotype in the *n*th dimension of individual *i* is Φ_*i*,*n*_ + 𝒩 (0, 1). In other words, we measure phenotypes on the natural scale set by mutation.

Strategies are controlled by three loci. The first locus is a switch whose two alleles toggle the active strategy type between innate and plastic. We refer to mutations to this locus as “inter-strategy” mutations. The second locus control the magnitude of the innate threshold *T*_*i*_ and the third locus controls the magnitude of the plastic threshold coefficient *a*_*i*_. We refer to mutations to these loci as “intra-strategy” mutations. At each of the three loci, mutations occur with probability *μ*_*S*_ = 1*/N* per individual per generation. For inter-strategy mutations, a mutation refers to a simple switch of their active strategy type. For intra-strategy mutations, new thresholds and threshold coefficients are drawn randomly from a uniform range of values, i.e., a house-of-cards mutation model. Note that even when a strategy type is not actively being used by the individual, its associated threshold or threshold coefficient is still evolving under drift and accumulated mutations.

## Results

### Natural selection favors plastic strategies in evolving populations

Our simulations show that populations consistently evolve to be primarily composed of plastic rather than innate strategists (Figure 2). This result is robust to changes in population size, mutation rate, payoff values, phenotype dimensions and initial conditions (See Appendix Figure 6). Figure 2 depicts a typical invasion (parameters listed in Table 1). Once plastic strategies have successfully invaded, they are evolutionarily stable, with coefficients remaining within a narrow range around *a*_*∗*_ ≈ 0.8. Levels of altruism among plastic strategists are substantially decreased from that of innate populations but above that of obligate defection. Innate populations on average donate ≈ 2 times per every 10 interactions, whereas plastic populations donate ≈ 1 time per every 20 interactions.

**Figure 2:**
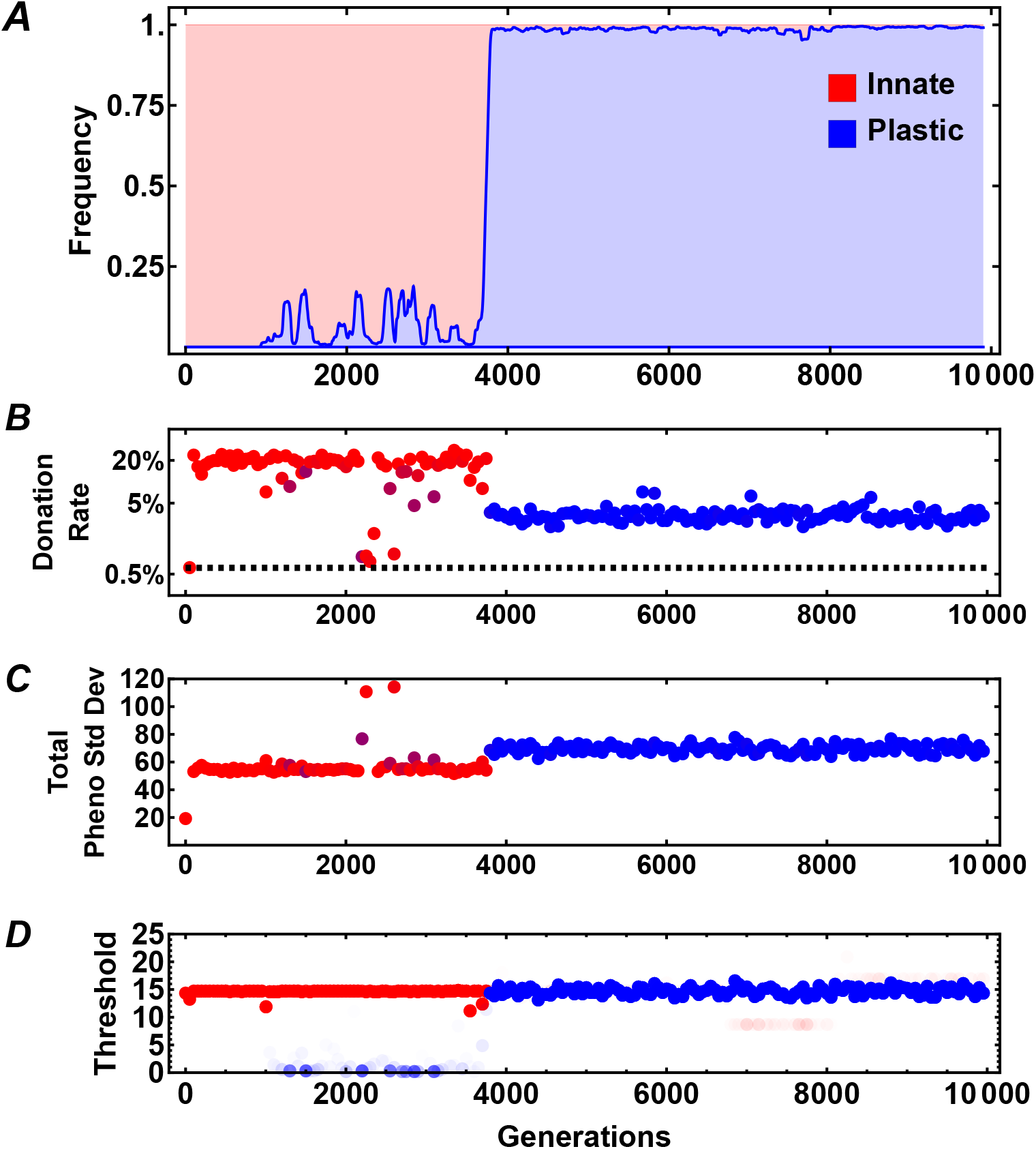
Plastic strategies are usually evolutionarily favored over innate strategies. Plots show the trajectory of a typical evolutionary simulation, with the horizontal axis being time in generations. We initialize the population with only innate strategies and allow it to equilibrate for 1000 generations before introducing inter-strategy mutations at a rate *μ*_*S*_ = 1*/N*. The vertical axis in each plot shows: **(A)** The frequency of plastic strategies (blue). Plastic strategies invade and remain resident. **(B)** Donation rate, with color indicating the frequency of innate (red) or plastic (blue) individuals. Donations drop when plastics invade. However, the population remains altruistic, in that donation is *≈* 10*×* than it would be if altruistic actions had no effect on selection (black dashed line). **(C)** The average total standard deviation in phenotype space of the population. Populations using the innate strategy form more dense clusters in phenotype space. **(D)** The average threshold used by the population. Both innate and plastic populations use the same average threshold, which is the expected stable threshold according to previous literature [Båvik et al., 2023]. While in innate populations, the average threshold is an accurate reflection of the real threshold used by most individuals, in plastic populations individuals with more close neighbors will use a smaller threshold than the depicted average, while individuals with rarer phenotypes will use larger thresholds than the depicted average threshold.

### Plastic strategists approximate the optimal threshold given their position in the population

In this section, we derive an expression for the optimal threshold that any individual in our simulations should use, given its genotype and phenotype and those of the rest of the population. This optimum is the donation threshold *T* that would maximize the expected change in frequency of the focal individual’s strategy allele in that generation. Intuitively, this is the threshold that maximizes the number of individuals sharing the strategy allele that are included while minimizing the number of individuals with different alleles that are included.

In our simulations, a focal individual chooses whether to donate to individuals within a sample *K* potential recipients based on their phenotypes. Let *p* be the frequency of the focal individual’s strategy allele, and let the prefix Δ denote the change in quantities due to the donations. We wish to find the threshold *T* that maximizes Δ*p*, the expected change in the frequency of the focal strategy allele. Label all individuals in the population 0 through *N* −1, with 0 labeling the focal individual, 1 through *K* labeling the potential recipients in increasing order of phenotypic distance to 1, and *K* + 1 through *N* labeling the remaining individuals. Then 0’s threshold *T* will determine how many recipients there are, which we will write as *Ñ* (*T*). Individuals with indices 1 through *Ñ* will receive donations, and others will not. Since the potential recipients are at *K* discrete positions in phenotype space, Δ*p* will be maximized by a range of thresholds that all give the same maximally distant recipient *Ñ*. If, for instance, the four closest potential recipients share the focal individual’s strategy allele and the six farther potential recipients do not, any threshold between *d*_04_ and *d*_05_ will result in *Ñ* = 4 and will be optimal. If all the *K* potential partners share the allele, then *Ñ* = *K* and the threshold could be arbitrarily large while still being optimal.

Let *W*_*i*_ be the payoff of individual *i* prior to 0’s donations. The payoffs then change by Δ*W*_0_ = − *Ñ c* and Δ*W*_*i*_ = *b* for 1 Δ *i* Δ *Ñ*, with other individuals’ payoffs unchanged, Δ*W*_*i*_ = 0 for *i > Ñ*. All individuals’ fitnesses *w* are affected because the mean payoff is increased by 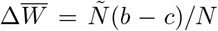, reducing fitness via the normalization in Eq. 1. To find the effect Δ*p* on the focal allele frequency due to the donation (or, equivalently, the effect *N* Δ*p* on the number of individuals carrying the focal allele), we need to weight the changes in each individual’s fitness by whether or not they are carrying the allele. This can be written as 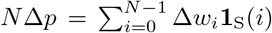, where **1**_S_(*i*) is an indicator function that is 1 if *i* carries the focal allele and 0 otherwise. (This implies that 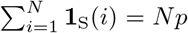.) The total expected change in the focal allele frequency is then:

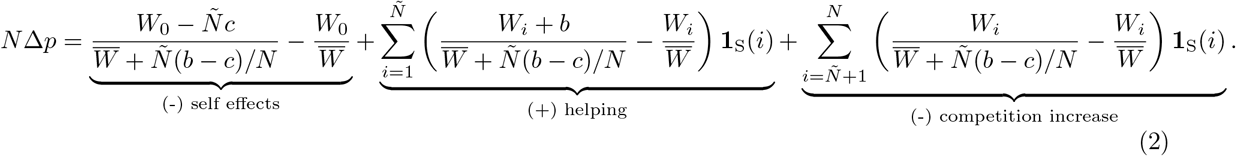

The optimal range of thresholds *T* is the one for which *Ñ* (*T*) maximizes Eq. 2. Figure 3 shows the accumulated value of Eq. 2 in simulated populations over a range of average distances from the focal individual to its partners (horizontal axis) and hypothetical altruism thresholds for the focal individual (vertical axis). The plastic altruists evolve to use thresholds that are close to optimal for their position in phenotype space. In contrast, the context-independent strategy used by innate altruists yields thresholds that are far from optimal given their mean partner distance and even disadvantageous relative to doing nothing. Recall that both the plastic and innate strategies have a single free parameter, so it is not that the plastic strategy can fit a wider range of optimal threshold patterns, but rather that its functional form is a better match to the typical optimal threshold pattern.

**Figure 3:**
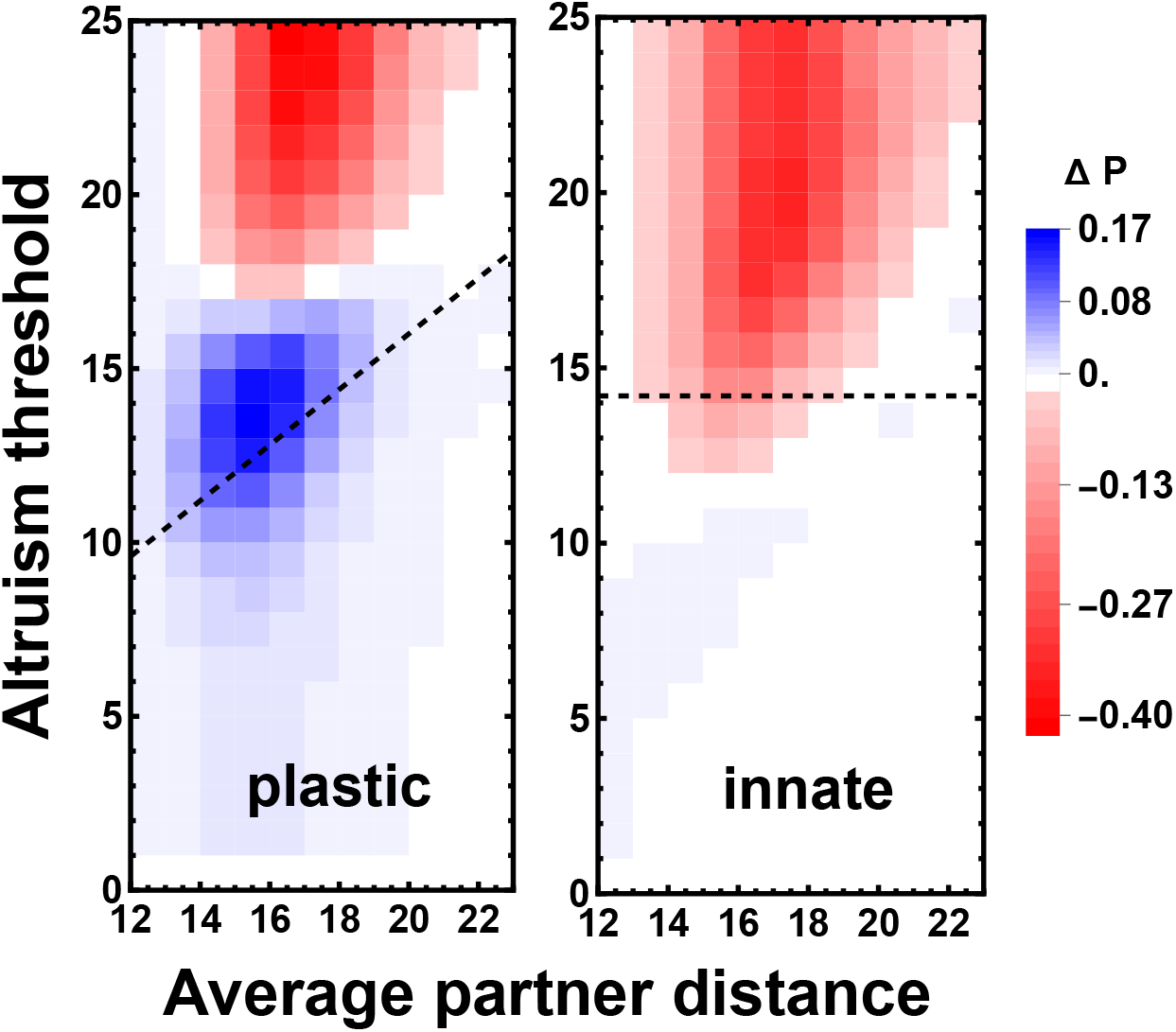
Plastic altruists obtain thresholds that yield positive strategy fitness at a range of phenotypic locations during invasion of an innate population. Plot is based on data from five replicate simulated populations of *N* = 1000 individuals. Each started as all innate altruists with the evolutionarily stable altruism threshold *T* = 14.2. After allowing the phenotype distribution to equilibrate for 10000 generations, we turn on strategy mutations to and from plastic altruists with threshold coefficient *a* = 0.8, who proceed to invade and take over. The horizontal axes of the plots show focal individuals’ observed mean phenotypic distances ⟨*d*_*i*_⟩ to their *K* = 10 potential recipients. The vertical axes show hypothetical altruism thresholds *T* that could be used by the focal individual, with the true thresholds that they used indicated as dashed lines. The left panel shows the results taking plastic altruists as the focal individuals, while the right panel takes innate altruists as the focal individuals. The color of each square with coordinates (⟨*d*_*i*_⟩ , *T*) shows by how much the panel’s strategy allele would have increased in frequency over the course of the simulation if individuals with the allele at mean distance ⟨*d*_*i*_⟩ had always used threshold *T*. This is calculated by summing Eq. 2 over all such individuals over the course of the simulations during the periods when the plastic strategy was at intermediate frequency, between 0.15 and 0.85. (Thus the values of Δ*p* along the dashed lines marking the true thresholds sum to the true allele frequency change, ±0.7.) The color of square therefore reflects both the payoff of an individual at ⟨*d*_*i*_⟩ using threshold *T* and the frequency with which individuals find themselves at ⟨*d*_*i*_⟩. The optimal (bluest) threshold tends to increase as a function of mean partner distance ⟨*d*_*i*_⟩, a behavior that the plastic strategy can match better than the innate one. Note that the plastic strategy is still suboptimal though, being slightly too selfish at typical mean partner distances of ⟨*d*_*i*_⟩ ≈ 14 − 17.

Eq. 2 provides an exact expression for the effect of one individual’s donations on the expected change in their strategy allele frequency, but it is hard to interpret. In our simulations, because the maximum number of recipients *K* is small compared to the population size *N* , the focal individual’s donations have only a small effect on the mean payoff of the population,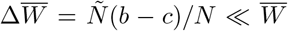. In this case, we can Taylor expand the denominators in Eq. 2, yielding a simpler approximate expression:

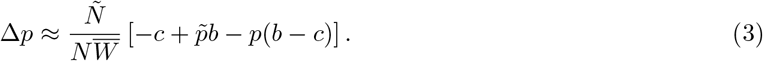

Here 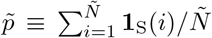 is the fraction of recipients who carry the focal allele, i.e., the average identity between the donor and recipients. The total effect is approximately just the sum of *Ñ* independent donations. For each one, the term in brackets is similar to the −*c* + *rb* that appears in Hamilton’s rule. (More precisely, we have 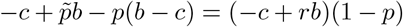, with 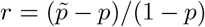 being the average relatedness of the recipients to the donor.)

### Plastic altruists outcompete innate altruists regardless of initial conditions

To support the results shown in Figures 2 and 3, we conduct simulations of the evolutionary dynamics of different innate and plastic strategies in a pairwise fashion in Figure 4. In these pairwise simulations, the value of the available plastic threshold coefficient, *a*, and the value of the available innate threshold, *T* , are fixed, so that all mutants between strategy types (plastic-to-innate or vice versa) must use the selected coefficient *a* if plastic or *T* if innate. This way, we can observe a “snapshot” of the evolutionary pressures on a population with a given pair of strategies, and how these pressures may influence the trajectory of populations through strategy space.

**Figure 4:**
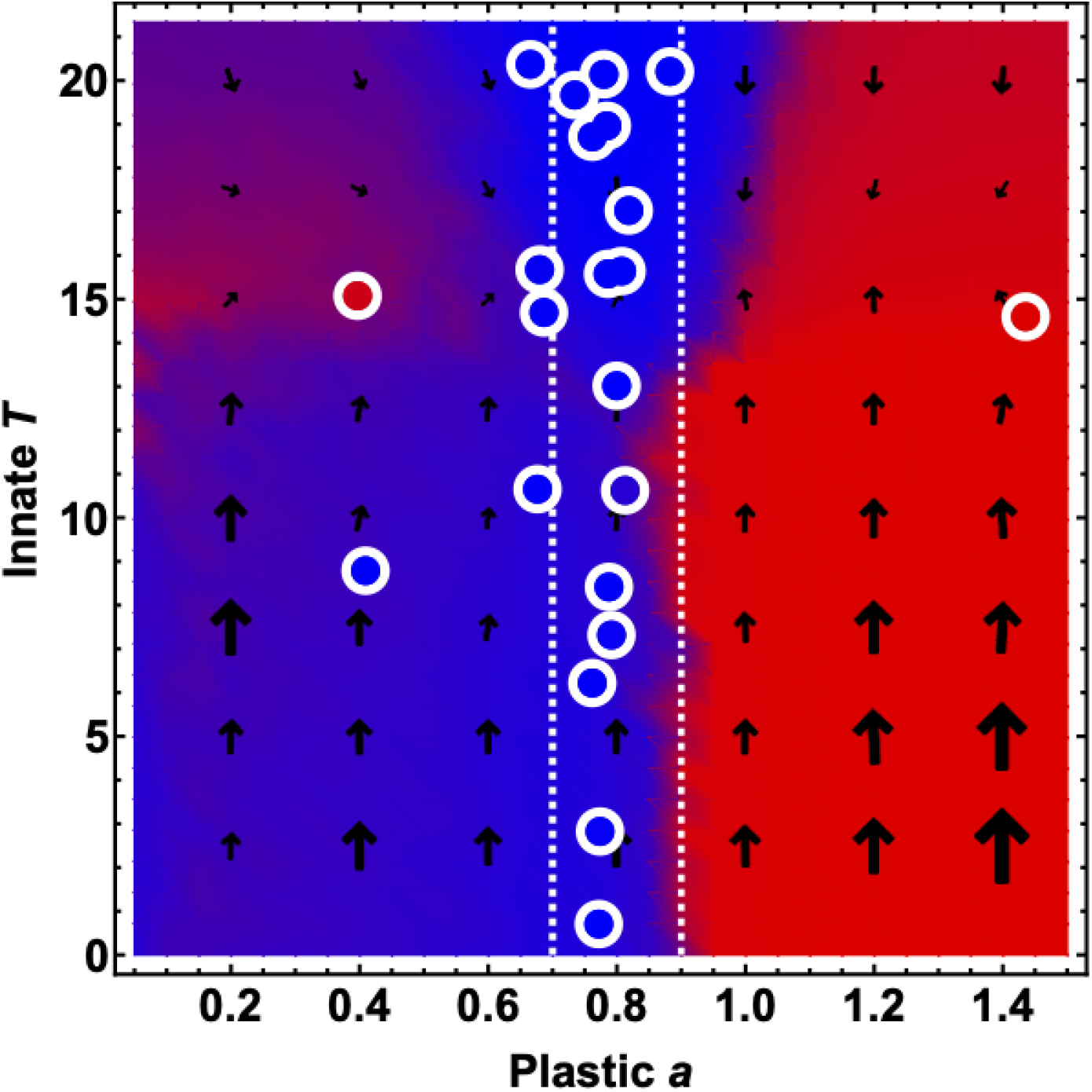
Plastic altruism using coefficient *a*_*∗*_ is the only strategy that can both invade and resist invasion by other strategies. We conduct a set of pairwise simulations of individuals with fixed thresholds. Fixed thresholds in this context means that mutations to threshold size within a strategy—intra-strategy mutations—are disabled, but mutations between strategy types—plastic-to-innate or innate-to-plastic, inter-strategy mutations—are allowed. Individuals evolve with resident strategy for 1000 generations, then inter-strategy mutation is introduced via a mutation rate *μ*_*S*_ = 1*/N*. The final frequency (averaged over the last 2000 out of 5000 generations) of the plastic strategy type is displayed in color that forms the background of the plot. Remarkably, the fixed threshold simulations show no effect from which strategy is resident and which is introduced via mutation, indicating no effect of initial conditions on inter-strategy competition. Finally we superimpose results of simulations allowing intra-strategy mutation (mutations to the magnitude of the plastic and innate thresholds, as was depicted in Figure 2) as the points with white edges. We run 25 simulations with initial conditions distributed evenly across the plane for 50,000 generations. Final coordinates of trajectories are displayed as points, colored by final frequency of plastic strategy. Vector arrows are calculated from the average change in threshold and threshold coefficient throughout the simulations (arrow size proportional to magnitude).

Figure 4 confirms that the evolutionarily stable state is a population dominated by plastic individualswith threshold coefficient *a* ≈ *a*_*∗*_. This result holds regardless of whether the populations are initially fixed for innate or plastic individuals (Appendix Figure 7). However, larger values of *a* are disfavored against a wide range of innate thresholds. Intuitively, *a* ≈ 1 corresponds to helping the average observed individual and serves as an upper bound on positive kin selection. Smaller values of *a* are generally successful against innate thresholds, but are disfavored against innate threshold *T* ≈ 14.2, which we previously found be the evolutionarily stable threshold for purely innate populations with the same values of the other parameters [Båvik et al., 2023].

### The role of sensing noise in context-dependent strategies

Above, we found that simulated populations that mutate between innate and plastic strategies tend to converge on a plastic strategy with coefficient *a*_*∗*_. One might expect that since innate individuals are very rare, one could simply remove the possibility of innate strategies from the simulations, and the resulting purely plastic populations would evolve to the same coefficient *a*_*∗*_. However, when we conduct these simulations we instead see something more complicated: the population’s strategy alternates between *a*_*∗*_ and a much lower threshold coefficient of order 10^*−*3^ *< a <* 10^*−*2^.

Figure 5C shows the distribution of strategies during plastic-only simulations. The dynamics appear to be bistable. One meta-stable state is the same one as found in the simulations with plastic individuals, with *a* ≈ *a*_*∗*_. In this case, the population forms a single cluster in phenotype space (Figure 5B), as we have observed previously in simulations of innate strategies [Båvik et al., 2023]. But another common state exists with much smaller *a*, corresponding to populations that have split into two widely separated clusters in phenotype space, with approximately equal numbers of individuals in each cluster (Figure 5A). Because the distance *δ* between the clusters in phenotype space is much larger than the cluster radii, individual *i*’s observed mean distance ⟨*d*_*i*_⟩ to its *K* potential partners is essentially determined by the number *n*_*i*_ of them it samples from the other cluster: ⟨*d*_*i*_⟩ ≈ *n*_*i*_*δ/K*—the precise positions of the individuals within their clusters only contribute small corrections to this. In other words, the observation ⟨*d*_*i*_⟩ only provides information about the irrelevant random number *n*_*i*_, not the focal individual’s place in the population’s phenotype distribution. In addition, *n*_*i*_ is drawn from a binomial distribution with only *K* draws: *n*_*i*_ ∼ Binomial(*K*, 1*/*2). This distribution has a substantial standard deviation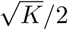 , which is ≈ 1.6 for *K* = 10, ≈ 30% of its mean value of 5. This means that plastic individual’s thresholds will vary widely (Figure 5D). Thus, even if *a* evolves to optimize the mean value of ⟨*d*_*i*_⟩, each individual’s value of ⟨*d*_*i*_⟩ will be sub-optimal due to fluctuations in its observations. Innate strategies do not have this disadvantage, and therefore invade when they are included in simulations (red strip at left edge of Figure 4.) These innate strategists are then rapidly replaced by plastic strategists with *a* ≈ *a*_*∗*_. This explains how innate strategies have a large impact on the simulations even though their average frequency over a simulation run is low.

**Figure 5:**
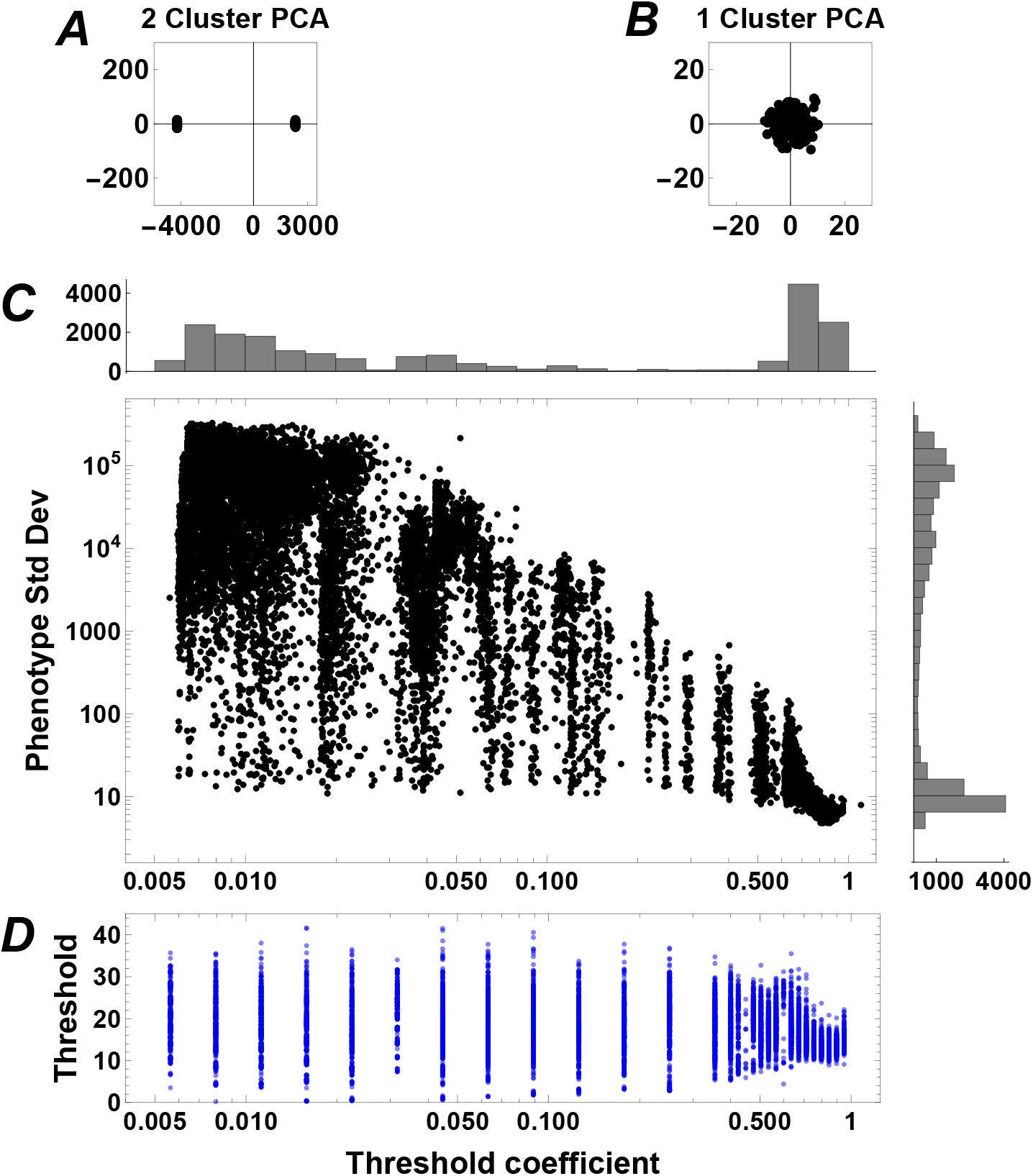
Evolution of a population of purely plastic individuals is bistable. **(A-C)** show the results of simulations in which strategies are restricted to be plastic (not innate) but the threshold coefficient *a* is allowed to evolve. The populations spend most of the simulations in one of two metastable states. **(A)** and **(B)** show the first two principal components of a population’s distribution in phenotype space in example generations when it is in the more and less frequent metastable states, respectively. **(A)**: The average threshold coefficient is *a* = 0.01, and population is split into two roughly equal clusters with a very large separation between them. **(B)**: The average threshold coefficient is *a* = 0.87 and the population comprises a single tight cluster. **(C)**: Populations are usually either in the two-cluster state with high phenotypic variation and threshold coefficient *a* ≪ 1, or the one-cluster state with low phenotypic variation and *a ≈ a*_*∗*_ *≈* 0.8. Points show the standard deviation of the population’s phenotypic tag against the average threshold coefficient over the course of 2 *×* 10^7^ generations of evolution (points taken at intervals of 10^4^). Histograms show the marginal distributions of the phenotypic standard deviation and the threshold coefficient *a*. **(D)** shows the range of thresholds used (blue points) in a vertical slice of the simulation in (C). Threshold data were binned into intervals (with smaller intervals used near *a ≈ a*_*∗*_ *≈* 0.8 to show the change in trend) and 1000 data points per bin were sampled to generate the data points. For small *a*, the population evolves to the two-cluster state and the fluctuations in thresholds used are on the order of average threshold. For large *a*, the population evolves to a single-cluster distribution with a smaller range of thresholds present.

**Figure 6:**
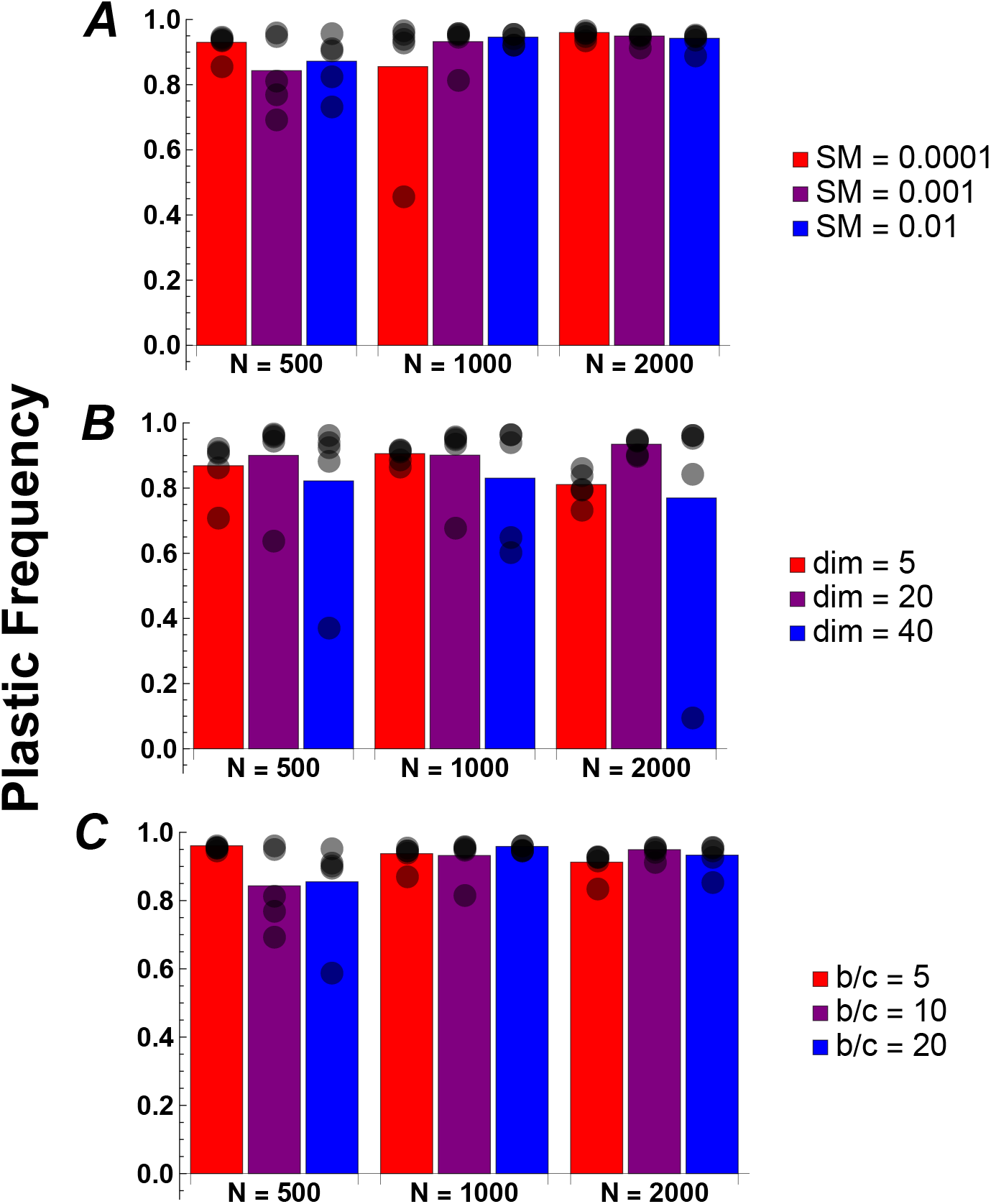
Evolution favors plastic strategies over parameter space. Plots show the frequency of plastic individuals in simulations allowing inter-strategy mutation. In each simulation run, the population was initialized as an innate population with thresholds at their typical end-state threshold (the cohesion threshold specified in Båvik et al. [2023]) and plastic threshold coefficients at *a* = 0.8. The population is initialized as innate individuals, and plastic individuals are introduced at a mutation rate 1*/N* after 1000 generations. Bars depict the frequency of plastic individuals across five iterations of simulations (each shown as a black dot) run for 50000 generations, and averaged over the last 5000. We vary the parameters used for simulations between a small (red), moderate (purple) and large (blue) value that is used for one set of simulations. Our simulations use the moderate (purple) values, and the moderate population size, *N* = 1000. In **(A)** we vary strategy mutation (SM), in **(B)** we vary phenotype dimension (dim) in **(C)** we vary the benefit-to-cost ratio (b/c). In most cases, plastic success is similar to the level we find using the parameters from Table 1.

**Figure 7:**
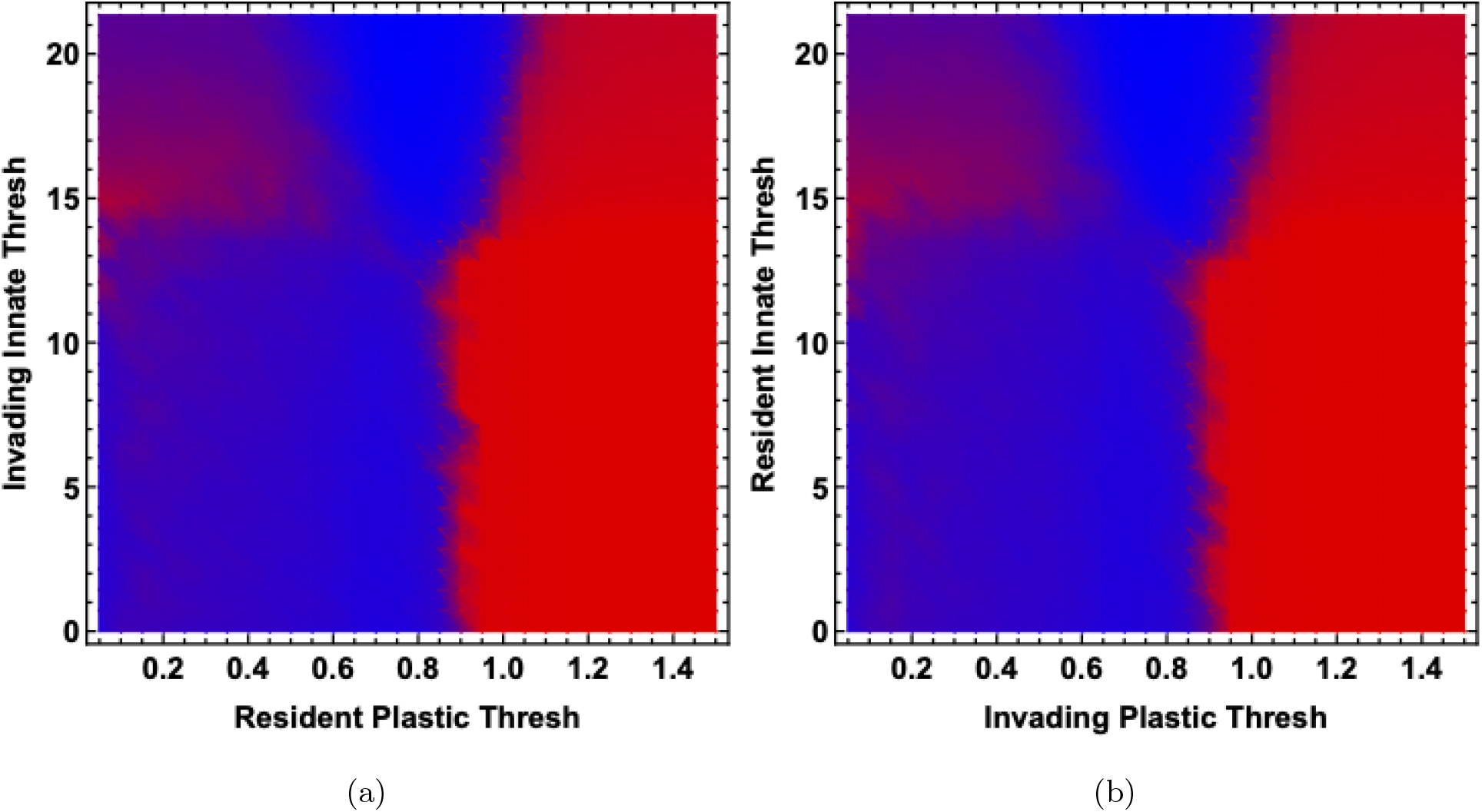
Residency conditions have no effect on the ultimate strategy frequency. This figure depicts two grids of pairwise invasion simulations of individuals with fixed thresholds in the manner described from Figure 4: Mutations to threshold size within a strategy—intra-strategy mutations—are disabled, but mutations between strategy types—plastic-to-innate or innate-to-plastic, inter-strategy mutations—are allowed. Individuals evolve with resident strategy for 1000 generations, then inter-strategy mutation is introduced via a mutation rate *μ*_*S*_ = 1*/N*. The final frequency (averaged over the last 2000 generations) of the plastic strategy type is displayed in color that forms the background of the plot. Remarkably, the fixed threshold simulations show no effect from which strategy is resident and which is introduced via mutation, indicating no effect of initial conditions on inter-strategy competition.

## Discussion

Our study supports the overall advantage of plasticity in altruistic decision making. The successful altruist often cannot simply measure dissimilarity between themselves and a potential partner: the decision to help or not should depend on who their other neighbors are. To summarize the logic: a population of innate strategists typically evolves to a single fixed threshold for altruism that is optimal for the individual who has neighbors at exactly the average distance. However, most individuals will either be closer or farther to their neighbors than this overall average. Hamilton’s rule dictates that these individuals should be stricter or more generous, respectively, but an innate threshold for altruism does not allow this. A plastic threshold allows individuals to come closer to matching Hamilton’s rule across the range of possible positions in the phenotype distribution. Our model thus suggests that Hamilton’s rule implies that plastic, context-sensitive altruism should typically be favored by natural selection. But our model also illustrates the importance of sources of noise to the optimal strategy. In some contexts, such as when the population is split into two distinct competing phenotypic clusters, an innate strategy is better than a plastic one. (This advantage is short-lived, however, as the invasion of innate strategists collapses the population back to a single cluster, in which plastic strategists are again favored.) In this case, plastic individuals have difficulty measuring their position within their cluster because of the noise introduced by the much larger divergence between clusters. Plasticity in altruism is therefore a likely but not certain outcome of evolution under our current theoretical understanding, highlighting the need to test whether it is in fact common in nature.

Our model has focused on a single well-mixed population, but population structure very likely has large effects on the evolution of altruism. Consider a population consisting of demes with occasional migration between them, with local density regulation so competition occurs within demes. Then an individual will typically be resident in a deme of relatives, but occasionally will be a migrate into a deme of more distantly related individuals. Intuitively, in the former case the focal individual should have a relatively strict threshold for helping, while in the latter case it should be more generous, as even an individual with a significantly different phenotype might have a substantial positive relatedness if the rest of the deme is even more different. Spatial structure thus might increase the range of situations that a lineage experiences through time, and thereby increase selection for plasticity. But this remains to be tested in an explicit model.

Beyond spatial structure, there are several important possible extensions to the simple model considered here. On the technical side, we have chosen a “switch” type model, in which individuals must choose either a purely plastic or purely innate strategy. This could be modified to a continuum between the two; individuals evolve a strategy that weights a hard-coded and context-dependent contribution into a final threshold. On the side of realism, we have presented a model in which individuals sense a population before making any altruistic decisions. This may be realistic for certain systems, such as nest or hive setting where an aggregate background odor is accessible. In other systems, organisms may need to make decisions on the fly, updating their population template in a partner-by-partner fashion. This tweak to the model could address the question, how much information do individuals truly need? Are there some cases where only a few partners need to be surveyed, or where it is better to keep updating one’s template continuously? Lastly, sexual reproduction and recombination could be considered. Signals in our model are composed of patterns that arise in the distance distributions in phenotype space. These patterns are likely to be substantially altered by sexual reproduction, which should smooth the phenotype distribution.

## Supporting information

fig7a_high_resolution

fig7a_high_resolution

---

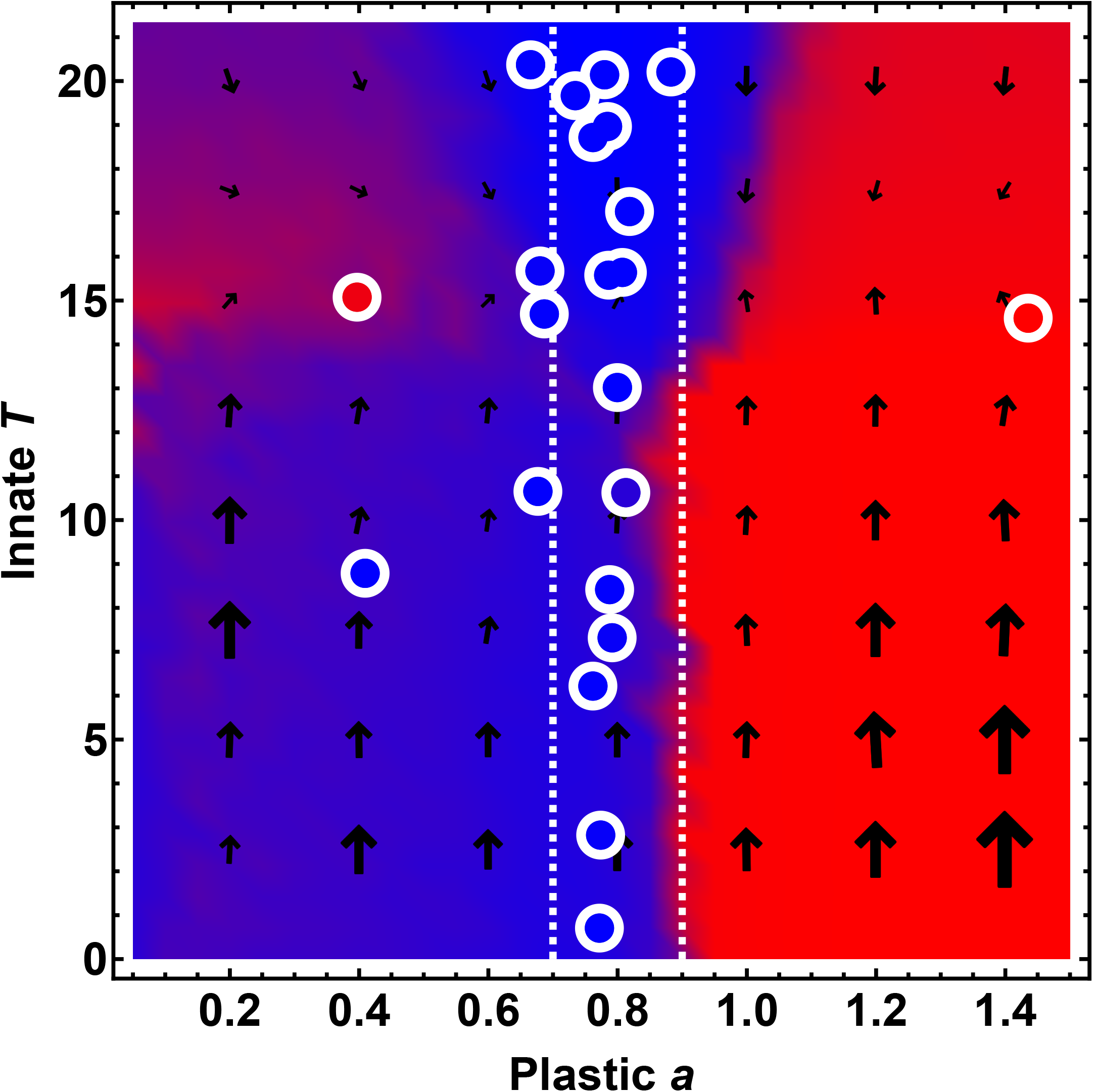

## Notes

### Competing Interest Statement

The authors have declared no competing interest.

