## Supplementary figures and images for "Selection favors context-dependent bias in altruism"

### fig7a_high_resolution

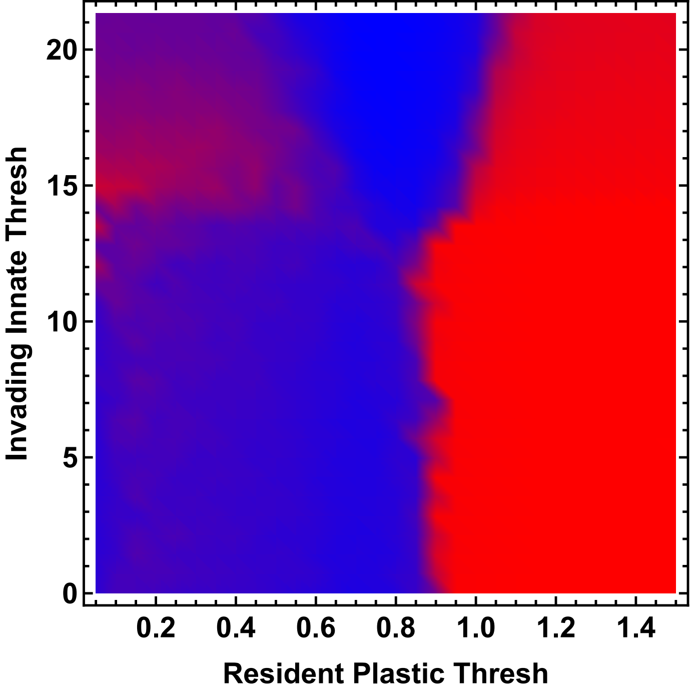

### fig7a_high_resolution

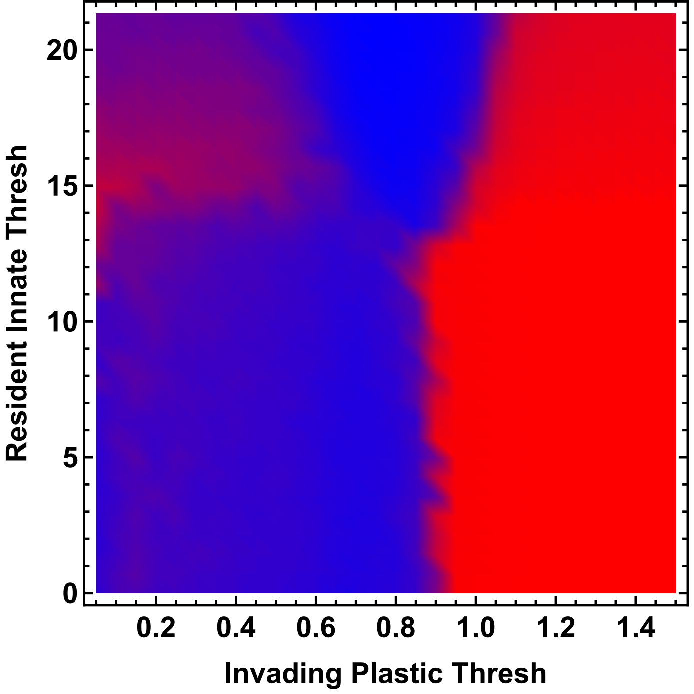
